# Structural and Functional NIR-II Fluorescence Bioimaging in Urinary System via Clinically Approved Dye Methylene Blue

**DOI:** 10.1101/2020.01.24.917955

**Authors:** Dingwei Xue, Di Wu, Zeyi Lu, Abudureheman Zebibula, Zhe Feng, Jun Qian, Gonghui Li

## Abstract

Accurate structural and functional imaging is vital for the diagnosis and prognosis of the urinary system diseases. Near-infrared region (NIR) II fluorescence imaging has shown advantages of high sensitivity, high safety, and fast feedback compared to the conventional imaging methods but limited to its clinical applicability. Herein, we first report that in vivo NIR-II fluorescence imaging of the urinary system enabled by clinically approved and renal-clearable NIR dye methylene blue, which can not only achieve clear invasive/non-invasive urography but also noninvasively detect renal function. These results demonstrate that MB assisted NIR-II fluorescence imaging holds great promise for invasive/noninvasive structural imaging of the urinary system clinically and investigation of renal function in animal models preclinically.

## 1. Introduction

Diagnosis and prognosis of urinary system diseases were based on imaging tests such as excretory urography, retrograde urography, urological computerized tomography, etc. However, those methods have disadvantages such as ray exposure, contrast allergy and poor effectiveness[1–5]. Besides, the need for sensitive, real-time and safe imaging methods increased along with the number of minimally-invasive procedures being performed.

Noninvasive analysis of renal function is also essential for accessing urinary system diseases, especially unilateral kidney diseases[6]. As the accurate evaluation of renal function demands real-time imaging of kidneys at high contrast and high temporal resolution, currently single-photon emission computed tomography (SPECT), magnetic resonance imaging (MRI) and positron emission tomography (PET) are the major tools for both clinical diagnosis and preclinical renal function studies[7–9]. Likewise, these methods were suffered from high cost, limited access, and potential radiation exposure risk. Therefore, safe, low-cost and sensitive renal functional imaging techniques are extremely desired for clinical and preclinical kidney research. Near-infrared fluorescence imaging, a promising biomedical imaging method, has shown superior properties in clinical translation owing to its high sensitivity, high temporal resolution and fast feedback, but is abstracted to limited penetration depth[10–13]. As imaging modality improved rapidly, the NIR-II window (1000 to 1700 nm) fluorescence bioimaging was verified to exhibit better spatial resolution, higher signal-to-background ratio (SBR) and deeper penetration depth compared to conventional NIR-I (780 to 900 nm) window in more and more studies [14–18]. To date, several kinds of NIR-II fluorescence probes including quantum dots (QDs) [17, 19, 20], carbon nanotubes[21–23], and rare-earth nanoparticles[24–26] have performed outstanding NIR-II fluorescence in whole-body and microscopic imaging. However, most of them confronted the same challenge in the process of clinical translation for their uncertain pharmacokinetic/toxicokinetic and drug metabolism[4, 27, 28]. Therefore, there is an urgent demand for NIR dyes which balances the advantages of NIR-II imaging and clinical applicability. So for, it was only a clinically approved NIR-I dye indocyanine green (ICG) that has shown potential in clinical NIR-II fluorescence imaging due to NIR-II emission tail [15, 16, 29].

Methylene blue (MB), a renal-clearable NIR-I dye approved by the U.S. Food & Drug Administration (FDA), has been extensively adopted for imaging-guided surgeries such as the identification of ureter, localization of insulinoma and normal pancreas, intraoperative detection of breast cancer[30–33]. Intriguingly, the excellent molar extinction coefficient (71,200 M^-1^ cm^-1^ at peak absorbance (665 nm)) and high quantum yield (3.8%)[30] made MB a candidate for clinical NIR-II fluorescence imaging. Whereas, the application of MB in NIR-II fluorescence imaging has not been reported yet.

Herein, we first successfully detected NIR-II emission tail of MB and compared tissue penetrating capability in the NIR-I window and NIR-II window by imaging capillary tubes filled with MB aqueous solution submerged in 1% Intralipid® solution in vitro. Subsequently, in vivo comparison of intravenous urography and retrograde urography in the NIR-I and NIR-II window were performed in the mouse models. Moreover, the NIR-II emission tail also made non-invasive renal function in the mouse model feasible. The aim of our study is to evaluate the validity and feasibility of applying MB to the structural and functional NIR-II bioimaging in the urinary system and provide a brand new potential translation of NIR-II imaging into clinical and preclinical applications.

## 2. Result and Discussion

### 2.1 MB optical characterization

MB, a small molecule NIR-I dye (chemical structure was shown in Fig. 1A), had an absorbance peak at ∼ 665 nm, and the emission spectrum of MB aqueous solution was recorded on a silicon (Si) and an indium gallium arsenide (InGaAs) detector based spectrometer, indicating that its fluorescence emission extended into NIR-II window (Fig. 1B-C). Beyond 1000 nm wavelength, the QY of MB in water was calculated as 0.2% (Fig S1), which is based on the reference of IR-26 in Dichloroethane (DCE)(0.5%, in the NIR-II region). Meanwhile, MB showed higher NIR-II fluorescence intensity under 623 nm LED excitation (0.1 mg/mL, 30 mW/cm^2^) than a reported renal-clearable NIR-II dye (CH-1055-PEG) under 793 nm laser excitation (0.1 mg/mL, 30 mW/cm^2^)(Fig S2). In addition, MB also showed an ideal photostability in water under a continuous 623 nm LED (80 mW/cm^2^) irradiation for 60 min with negligible fluorescence decay (Fig S3).

**Figure 1.**
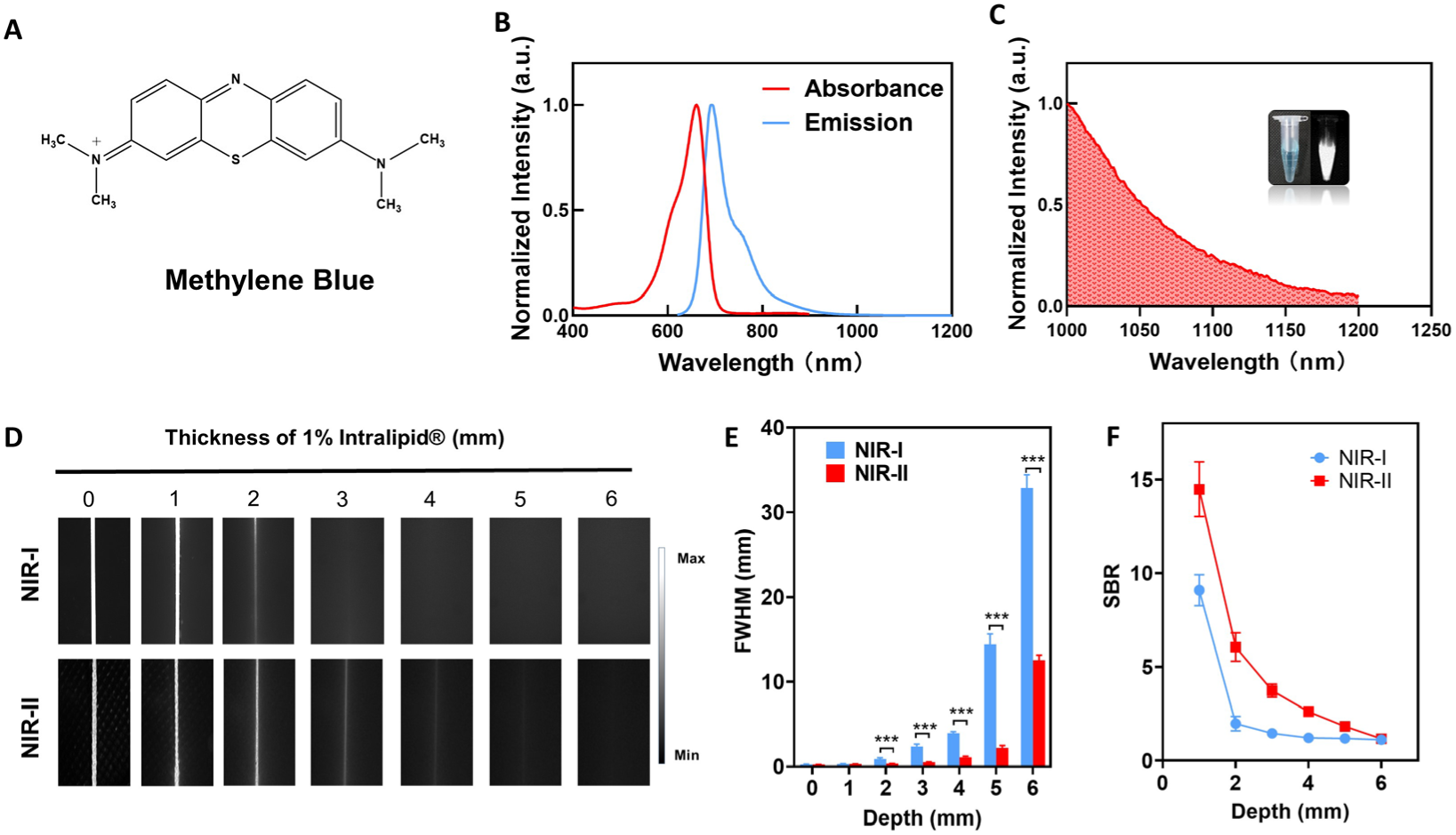
Characterization of MB. **(A)** Chemical structure of MB. **(B)** Normalized absorption and emission spectra of MB in water. **(C)** Normalized Fluorescence emission profile of MB between 1000 and 1200 nm wavelength region **(D)** NIR-I window and NIR-II window of glass capillary filled with MB (0.005 mg/mL) at depths of 0, 1, 2, 3, 4, 5, and 6 mm in 1% Intralipid® solution. **(E)** FWHM and **(F)** SBR were calculated for capillary glass tubes filled with MB solution. Data are the mean ± SD. n = 3 independent measurements.

To compare the penetrating ability of MB in NIR-I and NIR-II window, a tissue phantom study using Intralipid® mimicking the optical characteristics of biological tissues was performed. The fluorescence signals decreased both in NIR-I and NIR-II window with increasing thickness of 1% Intralipid® solution. The NIR-I fluorescence signal of MB was close to the background noise at a 3 mm thickness of 1% Intralipid® solution while the NIR-II fluorescence signal for MB was still visible even at a thickness of 5 mm. Full-width-half-maximum (FWHM) analysis depicting the feature width of NIR-I and NIR-II capillary images at varying depths in intralipid® phantom assay was also plotted (Fig. 1D): The FWHM measurement of the capillary tube without 1% Intralipid® solution were 386.5±5.4 μm and 389.4±1.7 μm in the NIR-I and NIR-II window, respectively. Whereas the FWHM was 3294.9±453.2 μm and 1243.1±14.4 μm in the NIR-I and NIR-II window, respectively, when the depth increased to 5 mm (Fig. 1E). In addition, the SBRs for MB in NIR-I window were 1.7, 4.3, 2.7, 2.1 and 1.8-fold higher than those for MB in NIR-II window at a 1% Intralipid® solution thickness of 1, 2, 3, 4, and 5 mm, respectively (Fig. 1F). These results indicated that MB fluorescence in the NIR-II window had deeper tissue penetration and higher sensitivity than those in the NIR-I window on account of the reduced light scattering in the NIR-II window.

### 2.2 Excreted behaviors study of MB

We initially conducted the excretion study of the MB. Fluorescence signals were mainly located in the bladder and gallbladder of mice injected with MB aqueous solutions (post-injection 30 min) in the NIR-II window (Figure S4), which demonstrated MB could be excreted by liver and kidney as reported in previous studies[34, 35]. Subsequently, the whole-body NIR-II fluorescence imaging (dorsal and ventral side) was conducted at different post-treatment time points after MB injection (Fig. 2A and 2B). The signal of the kidneys reached the maximum at 3 min post-injection of MB and then decreased with time (Fig. 2C), while fluorescence signals in the bladder increased with time (Fig. 2C). Furthermore, MB fluorescence exhibited minimal changes when diluted into mouse urine compared to MB aqueous solution at the same concentration (Fig. 2D and 2E). Taken together, MB showed a potential for real-time NIR-II fluorescence imaging of the urinary system including structural and functional imaging.

**Figure 2.**
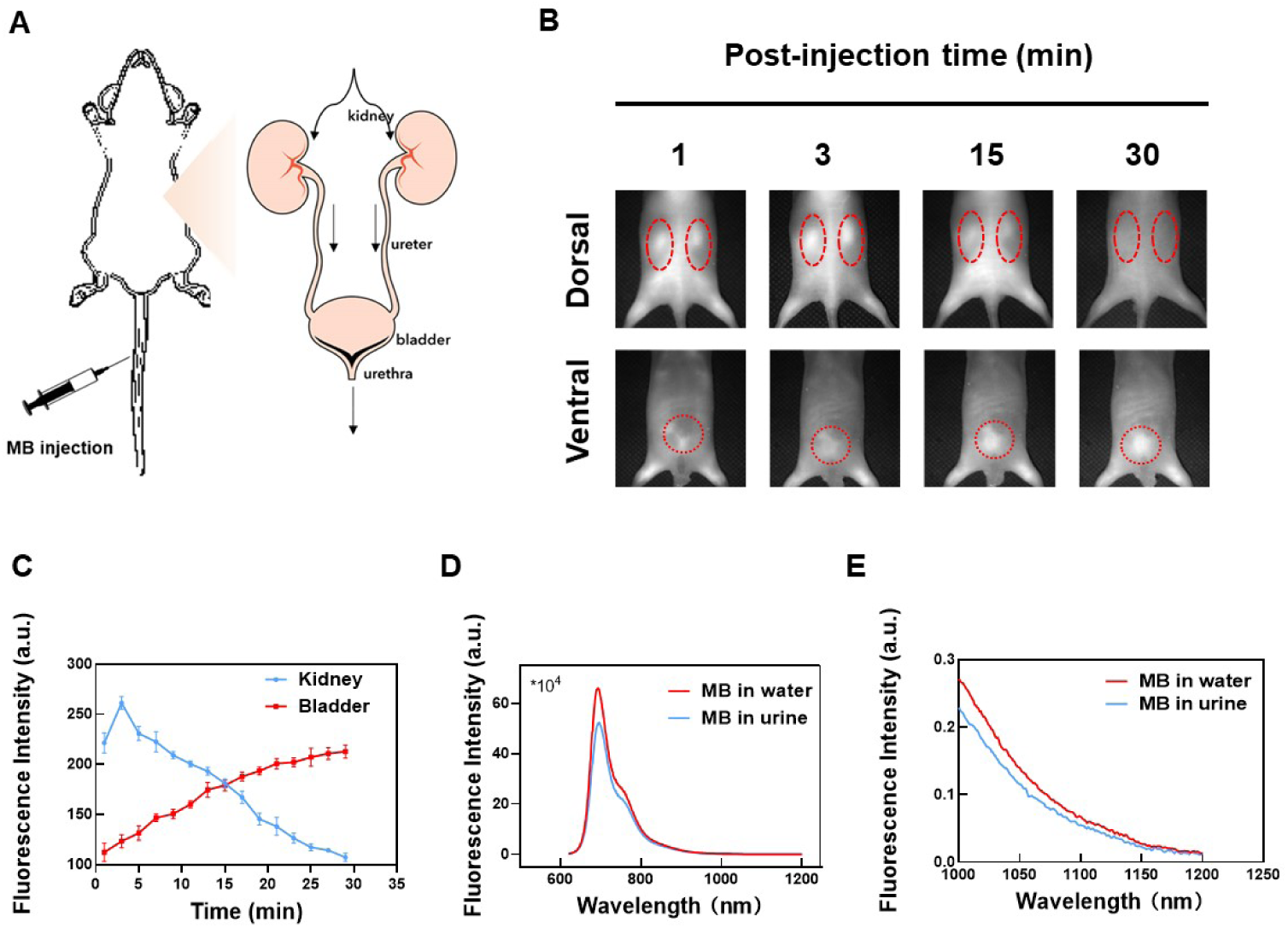
Renal clearance and in vivo stability studies of MB. **(A)** Schematic illustration of the renal excretion of MB through the urinary system. **(B)** Representative NIR-II fluorescence images at t =1, 3, 15, 30 min after injection of MB in mice. The red circles indicate the kidneys and bladder in the dorsal and ventral sides, respectively. NIR-II fluorescence images acquired beyond 1000 nm upon excitation at 623 nm, LED power ∼80 mW/cm^2^. **(C)** Time-NIR-II fluorescence intensity curves (TFICs) of the kidney and bladder after MB injection. Data are the mean ± SD. n = 3 independent measurements. **(D)** and **(E)** The emission spectrum of MB aqueous solution and MB in urine at the same concentration.

### 2.3 Structural imaging of the urinary system in vivo using MB in the NIR-I and NIR-II window

The NIR-II emission of MB enables a straightforward application to in vivo NIR-II fluorescence imaging. We first conducted non-invasive excretory and retrograde urography using NIR-I and NIR-II imaging in the same mouse model. It was obvious that imaging MB using NIR-II detection had advantages over conventional NIR-I detection: Imaging MB in the NIR-II window achieved high-contrast macroscopic imaging of kidney (excretory urography) and bladder (retrograde urography) in mice through intact skin (Fig. 3A and 3B).

**Figure 3.**
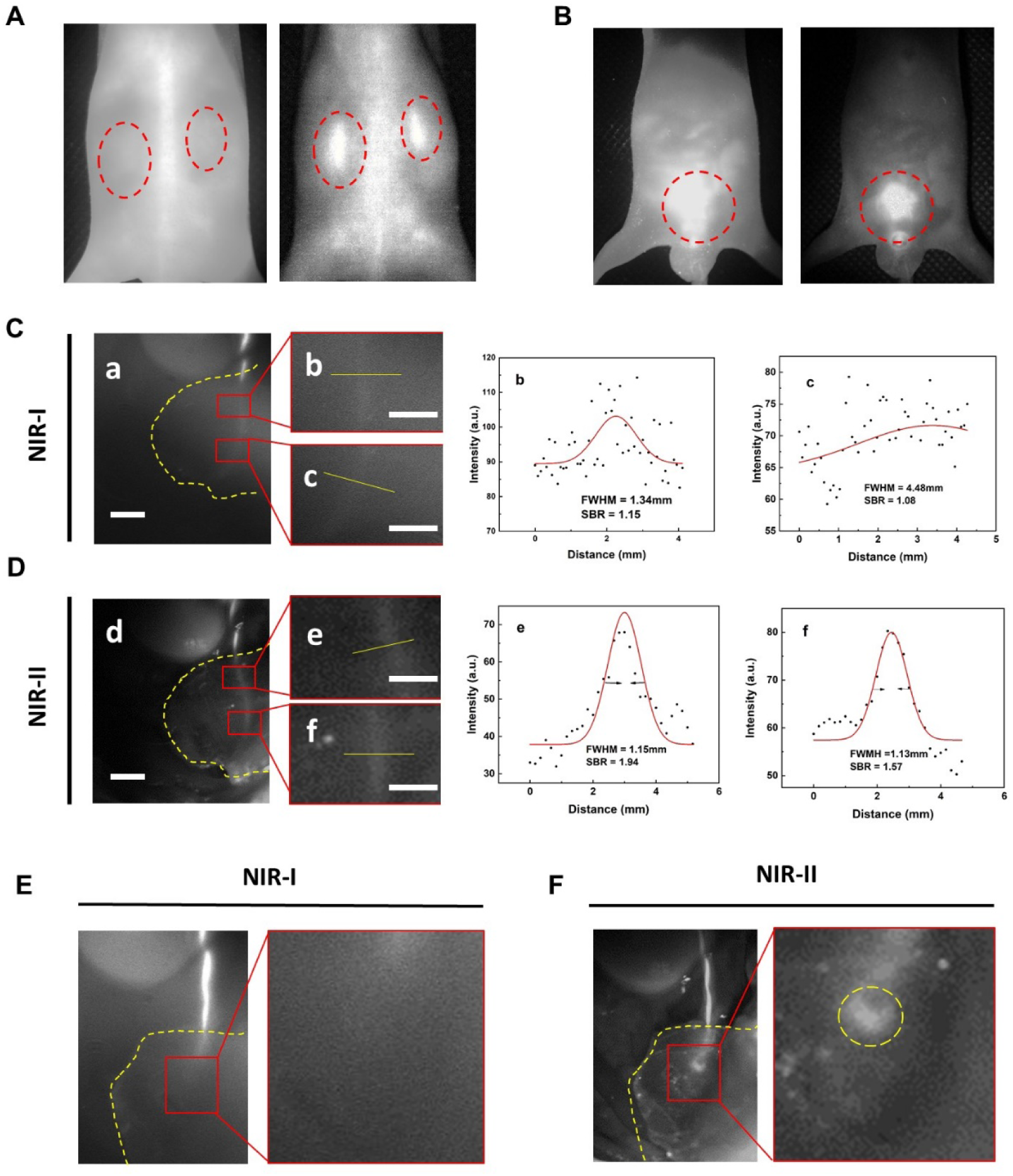
Imaging comparison of MB in NIR-I and NIR-II window in vivo. **(A)** Representative NIR-I and NIR-II fluorescence images of non-invasive images of the kidneys (0.01mg/1g body weight, i.v.) and **(B)** bladder (0.005mg/mL*20 μL, b.i.), respectively. **(C)** and **(D)** Representative images of ureter covered by mouse abdominal tissue after intravenous injection of MB in **(C)** NIR-I and **(D)** NIR-II window, FWHM and SBR of the ureter imaging in NIR-I and NIR-II window were calculated based on a two-term Gaussian fit to the intensity profiles. Scale bar in the left panel is 1cm; Scale bar in the right panel is 1mm. **(E)** and **(F)** Ureter imaging in the UUO model in the (E) NIR-I window and (F) NIR-II window, yellow circle showed the position of the obstruction point.

Ureter injury is a rare but serious complication of urinary surgery, however, the diagnosis of such injury is often delayed[36–38]. Thus, early identification of ureteral is essential to avoid morbidity and preservation of renal function. MB was reported to be successfully utilized in the intraoperative ureter visualization in the NIR-I window[31, 33]. We, therefore, detected the feasibility of NIR-II fluorescence imaging application in the ureter identification compared with the NIR-I window. Invasive NIR-I and NIR-II fluorescence imaging were applied to the real-time identification of ureter covered by a mouse abdominal tissue in the region of interest. Though the ureter could be visualized in the NIR-I window, the SBRs were rather low (Fig. 3C). When Switching to the NIR-II window with a 1000 nm long-pass filter, there was a significantly reduced background noise and improved spatial resolution (Fig. 3D). Whereafter, acute ureter obstruction was established on a mouse model to identify ureter lesion under NIR-I and NIR-II detector. As shown in Figure 3E and 3F, the ligation point of the ureter was obviously identified with good contrast in the NIR-II window while the specific ligation point could be hardly recognized under the NIR-I detector after upregulating the penetrating depth. These results indicated that MB assisted NIR-II fluorescence imaging was a superior choice in the image-guided surgery concerning ureter compared to the conventional NIR fluorescence imaging and could accelerate the clinical use of MB in fluorescence excretory and retrograde urography preoperatively and intraoperatively.

As full clinical implementation of MB has been partially limited when the ureter was covered by adhesion tissue image, the higher contrast of MB assisted NIR-II fluorescence imaging over NIR-I imaging could benefit these applications. Importantly, the implementation of this contrast improvement would be straightforward, requiring only a switch from cameras with NIR-I sensitivity to NIR-II-sensitive ones while continuing the familiar surgical setup and the use of MB.

### 2.4 In vivo MB Assisted NIR-II fluorescence imaging of renal function

Renal functional test or imaging usually reflects two aspects, that is, renal perfusion and renal filtration function. As the renal clearance is the main extracted way of MB and its fluorescence signal changes in kidneys could be detected non-invasively, MB seemed a good candidate for real-time imaging of renal function. Hence, the unilateral ureteral obstruction (UUO) model was first established for 3 d and 6 d (Fig. 4A) by complete ligation of the left ureter of the mouse while the right ureter was kept intact (Fig. 4B). As a result, unilateral hydronephrosis and renal perfusion disorder gradually progressed over time. For the sham-operated group, the ureters were not ligated. Subsequently, NIR-II fluorescence imaging was conducted in the UUO 3 d group, the UUO 6d group and the sham-operated group, and the profile for the signals in kidneys at different time points was recorded (Fig. 4C). With MB as the NIR-II contrast agent, we easily differentiated the UUO kidneys from the normal kidneys by noninvasive NIR-II imaging and analysis of the time-fluorescence intensity curves (TFICs) of the kidneys. As shown in Figure 4D-4F, the signal changes of the left kidney and right kidney showed no significant differences in the sham-operated group (Fig. 4D) while the obstructed left kidneys (LK) showed dramatically decreased signal peak value compared to the normal right kidneys (RK) in the UUO groups (Fig. 4G). Correspondingly, the peak time of the obstructed LK TFIC was delayed in the UUO groups compared to the normal RK in UUO groups and the kidneys in the sham-operated group (Fig. 4H). Moreover, the obstructed LK signals in the UUO 6d group showed a prolonged peak time compared to those in the UUO 3d group, which was consistent with the pathological analysis of kidney tissues (Fig. 4I): the renal tubules exhibited mild to moderate atrophy and dilatation in kidneys of the UUO 3 d group which suggested mild renal perfusion disorder, whereas, in kidneys in the UUO 6 d group, renal tubular damage and cortical atrophy were much more pronounced. These data indicated that MB assisted NIR-II fluorescence imaging of renal function could not only differentiate the normal kidneys and the kidneys with perfusion disorder but also reflect the severe stage of the perfusion disorder.

**Figure 4.**
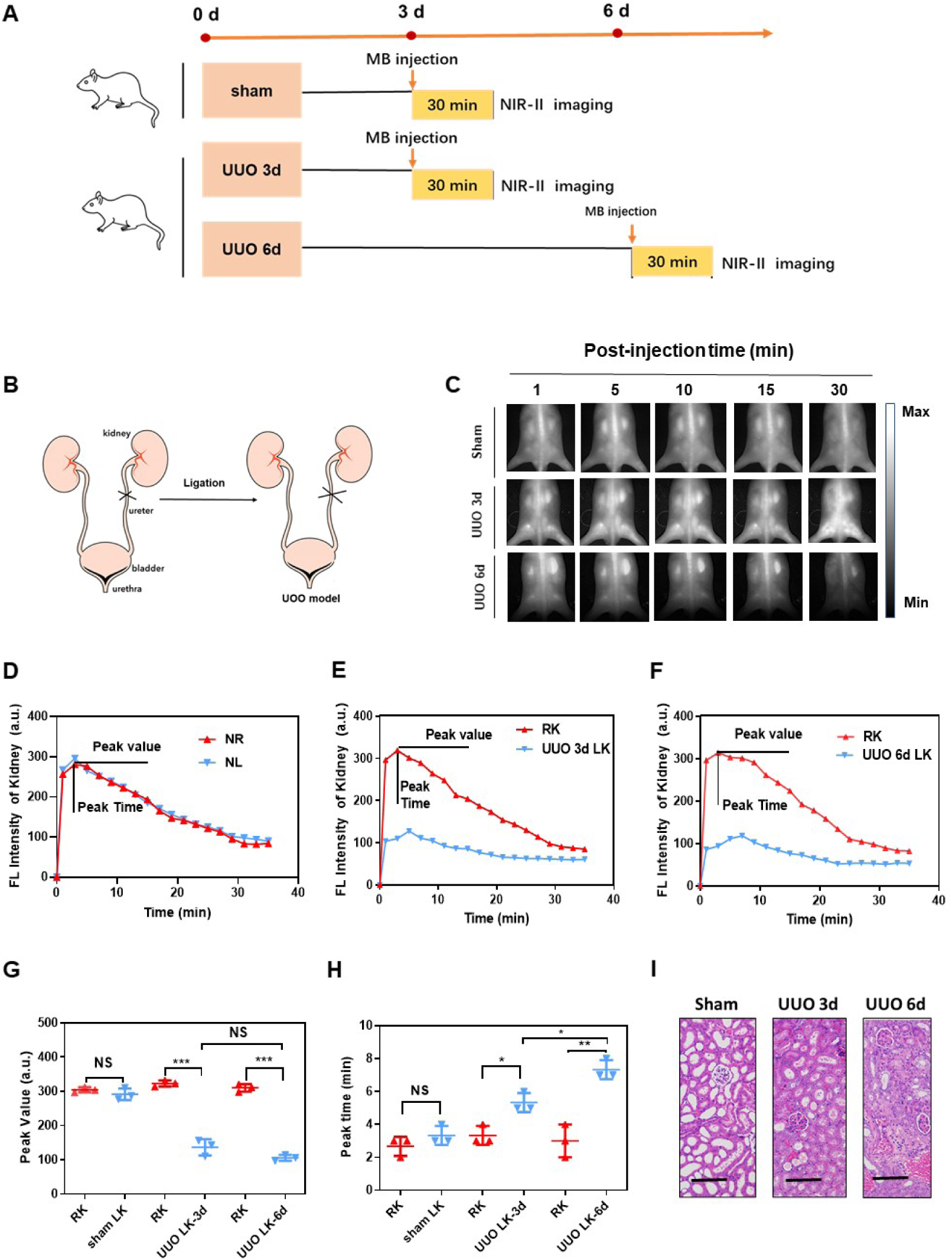
Real-time detection of kidney blood perfusion via NIR-II fluorescence imaging of MB in living mice. **(A)** Schematic illustration of mice of the sham-operated group and UUO group, and NIR-II fluorescence imaging at different post-operation timepoints. (**B)** Schematic illustration of UUO model establishment by complete ligation of the left ureter while the right ureter was kept intact. **(C)** Representative whole-body (dorsal side) noninvasive fluorescence images of mice after intravenous injection of MB solution (0.01mg/1g body weight, i.v.) at different post-injection time points. **(D-F)**. TFICs of kidneys in the sham control group, the UUO-3d group, and the UUO-6d group. **(G)** and **(H)** Statistical analysis of the two parameters extracted from the kidney TFICs of UUO mice and the sham control group. The parameters include the (G) peak value and (H) the peak time. Data are the mean ± SD. n = 3 independent measurements. *P<0.05, **P<0.01, ***P<0.001. (**I)** Kidney pathologic analysis of the sham-operated group, UUO-3d group, and UUO-6d group (H&E stain, scale bar=100 mm).

Renal filtration function is also indispensable for the renal function analysis, and blood urea nitrogen (BUN) and serum creatinine (Cre) are frequently used to evaluate the renal filtration function, but they both are not good indicators of a single kidney because of the presence of a well-functioning contralateral kidney. Given this compensatory mechanism, the solitary kidney model established by right nephrectomy was used to investigate the feasibility of MB assisted NIR-II real-time imaging in the assessment of renal filtration function. Further, the LKs were treated with vary degrees of electric coagulation injury in the unilateral renal failure group (URF) but were kept intact in the sham-operated group (Fig. 5A). Similarily, MB assisted NIR-II real-time imaging was carried out after MB i.v. injection with TFICs utilized to analyze the renal filtration function (defined as the clearance percentage at 30 min=[(peak value intensity at 30 min) /peak value]×100%)(Fig. 5B and 5C). As is shown in Figure 5D, MB clearance percentage at 30 min was reduced significantly in the URF group compared to the sham-operated group. To compare the renal function detection ability of MB with the clinical methods, Cre and BUN in the blood of living mice were measured using the commercial assays in the sham-operated group and the URF group. The statistically significant increases both in Cre and BUN were observed in the URF group, which was 1.67-fold and 1.38-fold higher than the sham-operated group, respectively. These data were consistent with the above imaging results, suggesting MB is feasible of noninvasively detecting renal filtration function by NIR-II fluorescence imaging.

**Figure 5.**
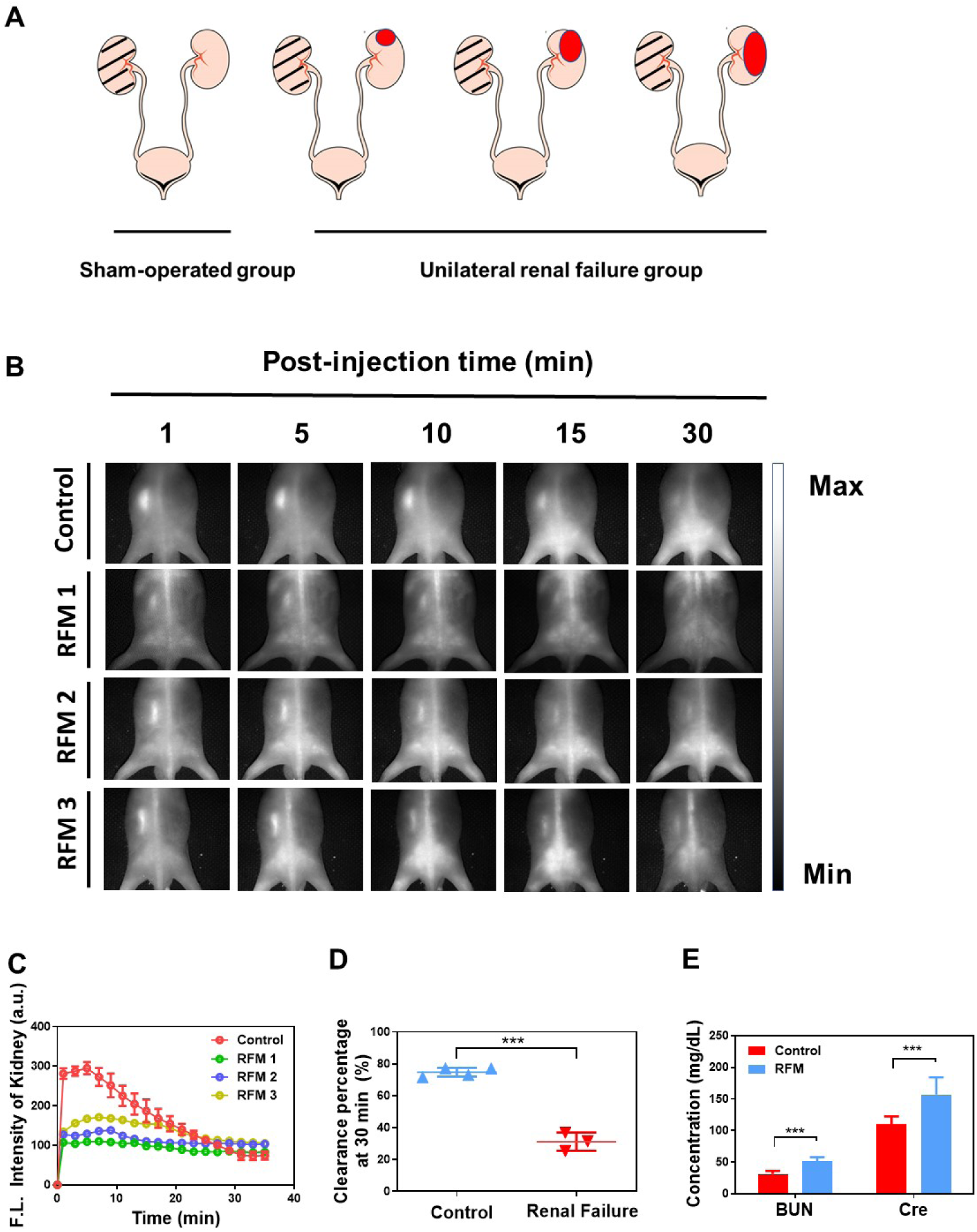
Real-time detection of renal filtration function via NIR-II fluorescence imaging of MB in living mice. **(A)** Schematic illustration of URF model (RFM) establishment (**B)** Representative NIR-II fluorescence images of the control group and RFM group after intravenous injection of MB (0.01mg/1g body weight, i.v.) at different post-injection time points. **(C)** TFICs of kidneys in the control group and RFM group. **(D)** Statistical analysis of the clearance percentage at 30 min extracted from the kidney TFICs of the control group and the URF group, ***P<0.001. **(F)** In vitro detection of renal function with other assays (Cre and BUN) in the control group and the URF group, ***P<0.001.

Although several NIR-II dyes such as CH-1055-PEG, CDIR2, rare-earth nanoparticles, and gold nanoparticles were reported to show potential in the NIR-II imaging of renal function[39–43], they all confronted a long way to extend the application in vivo with uncertain toxicity and pharmacokinetics. Moreover, the power densities of the excitation laser were fairly high in real-time imaging even for the animal models (300 mW/cm^2^ for both CDIR2 and CH-1055-PEG)[42, 44]. Nevertheless, MB could achieve clear real-time imaging at a relatively low power density of 623 nm LED excitation (80 mW/ cm^2^). As the absorbance of MB at 665 nm was 1.85 fold higher than that of MB at 623 nm, clear MB real-time imaging may achieve at an even lower power density at 665 nm excitation.

## 3. Conclusion

In summary, we detected the NIR-II emission of MB and investigated its application in the NIR-II fluorescence invasive/noninvasive urography and noninvasive NIR-II imaging of renal function in the mouse models. MB showed higher SBR and better spatial solution in the NIR-II window than those in the NIR-I window, which suggested a more appropriate detecting window when using MB fluorescence imaging clinically: By switching the detection of traditional silicon-based cameras to emerging InGaAs cameras could promisingly improve the fluorescence imaging technique both preoperatively and intraoperatively. As MB is excreted mainly through the kidney, the renal function analysis of MB assisted NIR-II fluorescence imaging is consistent with the pathology result and clinical diagnostic parameters including Cre and BUN. Thus, MB assisted NIR-II fluorescence imaging not only holds great promise for invasive and noninvasive structural imaging of the urinary system clinically but also permits investigation of renal function in living animal models preclinically.

## 4. Experimental Section

### 4.1 Materials

MB was purchased from JUMPCAN Pharmaceutical Factory (Taixing, China), which was used clinically. Phosphate buffer saline (PBS) was obtained from Sinopharm Chemical Reagent Company (Hangzhou, China). 20% intralipid® was purchased from Baxter Healthcare Corporation (USA). Deionized (DI) water with a resistivity of 18.2 MΩ/cm was used in all experiments.

### 4.2 Absorption and fluorescence emission characterization

Measurement of absorption spectra of MB aqueous solution was obtained from 550-900 nm using a Shimadzu UV-2550 UV–vis–NIR scanning spectrophotometer. The fluorescence emission spectra of MB dilutions in water and urine in the NIR-II window were measured by a lab-built system based on a PG2000 spectrometer (Ideaoptics Instruments) and a 2000C spectrometer (Everuping Optics Corporation).

### 4.3 Quantum yield measurement

The quantum yield of MB aqueous solution was measured using a NIR-II dye IR-26 in DCE as a reference (QY ≈ 0.5%) [45]. A series of DCE solutions of IR-26 and MB aqueous solution with different optical density (OD) values were measured under 623nm excitation, and NIR-II fluorescence intensities were integrated beyond 1000nm. Two slopes of the straight lines describing the dependence of integrated NIR-II fluorescence intensity upon OD (one from the reference of IR-26 in DCE and the other from the MB aqueous solution) were obtained. The *QY* of the sample (*Q_2_*) was calculated by the equation as follows:

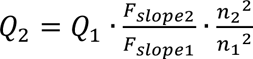

Where *Q_1_* is the *QY* of IR-26 in DCE (0.5%), *F_slope1_* is the slope value for IR-26 in DCE. *F_slope2_* is the slope value for MB aqueous solution. *n_2_* is the refractive index of water, *n_1_* is the refractive index of DCE.

### 4.4 The photostability assay

The photostability assay of MB aqueous solution (0.01mg/mL) was conducted under continuous illumination from 623 nm LED with a power density of 80 mW/cm^2^ for 60 minutes. The average fluorescence intensity was calculated from the region of the cuvette.

### 4.5 Intralipid® phantom imaging

In-vitro testing in an Intralipid® phantom was performed as described previously. 1% Intralipid® solution was prepared by diluting 20% Intralipid® into DI water. A capillary glass tube (Inner diameter = 0.3 mm) filled with MB solution (0.01mg/mL) was immersed in the prepared 1% Intralipid® solution, the depth of which ranged from 1-6 mm below the top surface. NIR-I and NIR-II imaging at different depths were performed.

### 4.6 Animal experiments

All the animal experiments in this work were conducted strictly in compliance with the requirements and guidelines of the Institutional Ethical Committee of Animal Experimentation of Zhejiang University. Institute of Cancer Research (ICR) mice (6-8 weeks old, female) and BLAB/c Nude mice (6-8 weeks old, female) were provided from the SLAC laboratory Animal Corporation. (Shanghai, China) and kept in the Laboratory Animal Center of Zhejiang University (Hangzhou, China). The animal housing area was maintained at 24 ℃ with a 12 hours light/dark cycle, with free water and food available.

Before each operation and imaging experiment, mice were anesthetized via intraperitoneal injection of 2% pentobarbital (40-50 mg/kg) and kept in maintaining anesthesia. Mice were intravenously injected with 200 μg MB before in vivo imaging.

### 4.7 NIR-I fluorescence imaging

Images in the NIR-I window were captured using GA1280 camera (1280 pixels ×1024 pixels, TEKWIN SYSTEM, China) equipped with a prime lens (focal length: 50 mm, antireflection coating at 800-2000 nm, Edmund Optics), which was fitted with an 800 nm long-pass filter and a 900 nm short-pass filter to extract NIR-I fluorescence signal. A 623 nm LED was used to provide uniform illumination on the field of interest.

### 4.8 NIR-II fluorescence imaging

A 2D electronic-cooling InGaAs camera (640 pixels×512 pixels, TEKWIN SYSTEM, China) equipped with a prime lens (focal length: 50 mm, antireflection coating at 800-2000 nm, Edmund Optics), cooled to −40 ℃ was used to acquire images in the NIR-II window (Fig. S5). A 623 nm LED (SOLIS-623C, Thorlabs, USA) was used to provide uniform illumination on the imaging field. The facular power density was measured before each imaging experiment. During imaging, an 800 nm long-pass filter was used to filter 623 nm excitation. A 1000 nm long-pass filter (ThorLabs, USA) was placed in front of the camera lens restricting wavelength below as well as allowing wavelength above 1000 nm to pass through the camera lens.

### 4.9 In vivo kidney and bladder structural imaging

Real-time NIR-II fluorescence imaging was conducted after i.v. injection of MB (10mg/kg body weight) in living mice and the signals of kidneys (dorsal side) and bladders (ventral side) were acquired beyond 1000nm (623 nm LED, 80 mW/cm^2^).

### 4.10 In vivo ureter imaging

Mice were fixed in a platform in the supine position, and laparotomy was performed and the ureters were fully exposed. 200 μg of MB was intravenously injected into each mouse, and then a piece of abdominal tissue was placed on the ureter. NIR-I and NIR-II imaging were further performed to visualize the ureter through the covered tissue. For the acute ureter ligation model, the right ureter was ligated by surgical sutures. Likewise, MB at a concentration of 10 mg/kg was intravenously injected through the tail vein instantly after ligation, and the facular power density was adjusted to 30 mW/cm^2^. The mice were observed using the NIR-I or NIR-II camera to localize the ureter and detect the ligation point covered by an abdominal tissue.

### 4.11 In vivo renal functional imaging

UUO model was established to investigate the renal perfusion by using MB assisted NIR-II fluorescence imaging. Briefly, the UUO model was first established by complete ligation of the left ureter of the mouse while the right ureter was kept intact (Fig. 4B), and unilateral hydronephrosis and renal perfusion disorder gradually progressed over time. For the sham-operated group, the ureters were not ligated. Subsequently, NIR-II fluorescence imaging was conducted in the UUO 3 d group, the UUO 6d group and the sham-operated group after MB i.v. injection (10mg/kg body weight). The profile for the signals in kidneys at different time points was recorded as a function of imaging time. On the other hand, the URF model was established to assess the renal filtration ability of MB assisted NIR-II fluorescence imaging. Briefly, the right kidneys were excised both in the URF model and the sham-operated group, and the left kidneys were injured by an electric coagulation knife to varing degrees in the URF model group and kept intact in the sham-operated group. NIR-II fluorescence imaging was conducted as mentioned above.

### 4.12 Serum creatinine and BUN assay

Blood was collected from the angular vein of the mice. The collected blood samples were centrifuged for 15 min at 4500 r.p.m. Serum creatinine and BUN were determined using commercial kits according to the manufacturer′s protocols.

### 4.13 Histopathologic study

The kidney tissues of the UUO groups and the sham-oparete group were dissected and fixed with 4% paraformaldehyde, dehydrated in an ethanol solution, embedded in paraffin and cut into sections with a thickness of 15 μm for H&E staining. The sections were washed with xylene and ethanol and then immersed in hematoxylin working solution for 4 min and eosin working solution for 2 min, followed by washing with distilled water. The stained sections were examined using a microscope (Primovert, Zeiss, Germany).

### 4.14 Data analysis

Quantitative analysis of each fluorescent image was performed based on the measurement of mean signal intensity in the manually selected regions of interest, using Image J software (Version 1.6.0, National Institutes of Health, USA). Graphs were generated using Origin Pro software (Version 9.0; OriginLab Company, USA). The data were presented as mean ± standard deviation (SD). Statistical analysis was performed using Student’s t-test. * denotes a statistical significance (* P< 0.05, ** P < 0.01, and *** P < 0.001) between the experiment data of two groups.

## Supporting Information

Supporting Information is available from the Wiley Online Library or from the author.

## Acknowledgment

This work was supported by the National Natural Science Foundation of China (Grant Numbers: 81672520, 81773789, 81870484, 81702508 and 61975172, used for procurement of materials and labor cost)

## Reference

1. Essman SC: Contrast cystography. Clin Tech Small Anim Pract 2005, 20(1):46–51.

2. McDonald RJ, McDonald JS, Carter RE, Hartman RP, Katzberg RW, Kallmes DF, Williamson EE: Intravenous contrast material exposure is not an independent risk factor for dialysis or mortality. Radiology 2014, 273(3):714–725.

3. Bjurlin MA, Turkbey B, Rosenkrantz AB, Gaur S, Choyke PL, Taneja SS: Imaging the High-risk Prostate Cancer Patient: Current and Future Approaches to Staging. Urology 2018, 116:3–12.

4. Moosavi B, Shabana WM, El-Khodary M, van der Pol CB, Flood TA, McInnes MD, Schieda N: Intracellular lipid in clear cell renal cell carcinoma tumor thrombus and metastases detected by chemical shift (in and opposed phase) MRI: radiologic-pathologic correlation. Acta radiologica (Stockholm, Sweden: 1987) 2016, 57(2):241–248.

5. Morris MJ, Autio KA, Basch EM, Danila DC, Larson S, Scher HI: Monitoring the clinical outcomes in advanced prostate cancer: what imaging modalities and other markers are reliable? Seminars in oncology 2013, 40(3):375–392.

6. Chawla LS, Eggers PW, Star RA, Kimmel PL: Acute kidney injury and chronic kidney disease as interconnected syndromes. The New England journal of medicine 2014, 371(1):58–66.

7. Taylor AT, Lipowska M, Cai H: 99mTc(CO)3(NTA) and 131I-OIH: comparable plasma clearances in patients with chronic kidney disease. J Nucl Med 2013, 54(4):578–584.

8. Grenier N, Basseau F, Ries M, Tyndal B, Jones R, Moonen C: Functional MRI of the kidney. Abdom Imaging 2003, 28(2):164–175.

9. Taylor AT: Radionuclides in nephrourology, part 1: Radiopharmaceuticals, quality control, and quantitative indices. J Nucl Med 2014, 55(4):608–615.

10. Cheng D, Peng J, Lv Y, Su D, Liu D, Chen M, Yuan L, Zhang X: De Novo Design of Chemical Stability Near-Infrared Molecular Probes for High-Fidelity Hepatotoxicity Evaluation In Vivo. J Am Chem Soc 2019, 141(15):6352–6361.

11. Feng Z, Yu X, Jiang M, Zhu L, Zhang Y, Yang W, Xi W, Li G, Qian J: Excretable IR-820 for in vivo NIR-II fluorescence cerebrovascular imaging and photothermal therapy of subcutaneous tumor. Theranostics 2019, 9(19):5706–5719.

12. Sun C, Li B, Zhao M, Wang S, Lei Z, Lu L, Zhang H, Feng L, Dou C, Yin D, Xu H, Cheng Y, Zhang F: J-Aggregates of Cyanine Dye for NIR-II in Vivo Dynamic Vascular Imaging beyond 1500 nm. Journal of the American Chemical Society 2019, 141(49):19221–19225.

13. Hori Y, Otomura N, Nishida A, Nishiura M, Umeno M, Suetake I, Kikuchi K: Synthetic-Molecule/Protein Hybrid Probe with Fluorogenic Switch for Live-Cell Imaging of DNA Methylation. Journal of the American Chemical Society 2018, 140(5):1686–1690.

14. Ding F, Zhan Y, Lu X, Sun Y: Recent advances in near-infrared II fluorophores for multifunctional biomedical imaging. Chemical Science 2018, 9(19):4370–4380.

15. Zhu S, Hu Z, Tian R, Yung BC, Yang Q, Zhao S, Kiesewetter DO, Niu G, Sun H, Antaris AL, Chen X: Repurposing Cyanine NIR-I Dyes Accelerates Clinical Translation of Near-Infrared-II (NIR-II) Bioimaging. Adv Mater 2018:e1802546.

16. Carr JA, Franke D, Caram JR, Perkinson CF, Saif M, Askoxylakis V, Datta M, Fukumura D, Jain RK, Bawendi MG, Bruns OT: Shortwave infrared fluorescence imaging with the clinically approved near-infrared dye indocyanine green. Proc Natl Acad Sci U S A 2018, 115(17):4465–4470.

17. Zebibula A, Alifu N, Xia L, Sun C, Yu X, Xue D, Liu L, Li G, Qian J: Ultrastable and Biocompatible NIR-II Quantum Dots for Functional Bioimaging. Advanced Functional Materials 2018, 28(9):1703451.

18. Del Rosal B, Villa I, Jaque D, Sanz-Rodriguez F: In vivo autofluorescence in the biological windows: the role of pigmentation. J Biophotonics 2016, 9(10):1059–1067.

19. Zhang M, Yue J, Cui R, Ma Z, Wan H, Wang F, Zhu S, Zhou Y, Kuang Y, Zhong Y, Pang DW, Dai H: Bright quantum dots emitting at approximately 1,600 nm in the NIR-IIb window for deep tissue fluorescence imaging. Proc Natl Acad Sci U S A 2018, 115(26):6590–6595.

20. Hong G, Robinson JT, Zhang Y, Diao S, Antaris AL, Wang Q, Dai H: In vivo fluorescence imaging with Ag2S quantum dots in the second near-infrared region. Angew Chem Int Ed Engl 2012, 51(39):9818–9821.

21. Diao S, Hong G, Robinson JT, Jiao L, Antaris AL, Wu JZ, Choi CL, Dai H: Chirality enriched (12,1) and (11,3) single-walled carbon nanotubes for biological imaging. J Am Chem Soc 2012, 134(41):16971–16974.

22. Robinson JT, Hong G, Liang Y, Zhang B, Yaghi OK, Dai H: In vivo fluorescence imaging in the second near-infrared window with long circulating carbon nanotubes capable of ultrahigh tumor uptake. J Am Chem Soc 2012, 134(25):10664–10669.

23. Hong G, Diao S, Chang J, Antaris AL, Chen C, Zhang B, Zhao S, Atochin DN, Huang PL, Andreasson KI, Kuo CJ, Dai H: Through-skull fluorescence imaging of the brain in a new near-infrared window. Nature photonics 2014, 8(9):723–730.

24. Wang R, Li X, Zhou L, Zhang F: Epitaxial seeded growth of rare-earth nanocrystals with efficient 800 nm near-infrared to 1525 nm short-wavelength infrared downconversion photoluminescence for in vivo bioimaging. Angew Chem Int Ed Engl 2014, 53(45):12086–12090.

25. Wang P, Fan Y, Lu L, Liu L, Fan L, Zhao M, Xie Y, Xu C, Zhang F: NIR-II nanoprobes in-vivo assembly to improve image-guided surgery for metastatic ovarian cancer. Nat Commun 2018, 9(1):2898.

26. Naczynski DJ, Tan MC, Zevon M, Wall B, Kohl J, Kulesa A, Chen S, Roth CM, Riman RE, Moghe PV: Rare-earth-doped biological composites as in vivo shortwave infrared reporters. Nat Commun 2013, 4:2199.

27. Alshehri R, Ilyas AM, Hasan A, Arnaout A, Ahmed F, Memic A: Carbon Nanotubes in Biomedical Applications: Factors, Mechanisms, and Remedies of Toxicity. Journal of medicinal chemistry 2016, 59(18):8149–8167.

28. Wang Y, Hu R, Lin G, Roy I, Yong KT: Functionalized quantum dots for biosensing and bioimaging and concerns on toxicity. ACS applied materials & interfaces 2013, 5(8):2786–2799.

29. Yu X, Feng Z, Cai Z, Jiang M, Xue D, Zhu L, Zhang Y, Liu J, Que B, Yang W, Xi W, Zhang D, Qian J, Li G: Deciphering of cerebrovasculatures via ICG-assisted NIR-II fluorescence microscopy. J Mater Chem B 2019, 7(42):6623–6629.

30. Winer JH, Choi HS, Gibbs-Strauss SL, Ashitate Y, Colson YL, Frangioni JV: Intraoperative Localization of Insulinoma and Normal Pancreas Using Invisible Near-Infrared Fluorescent Light. Annals of Surgical Oncology 2009, 17(4):1094–1100.

31. Verbeek FP, van der Vorst JR, Schaafsma BE, Swijnenburg RJ, Gaarenstroom KN, Elzevier HW, van de Velde CJ, Frangioni JV, Vahrmeijer AL: Intraoperative near infrared fluorescence guided identification of the ureters using low dose methylene blue: a first in human experience. J Urol 2013, 190(2):574–579.

32. Tummers QRJG, Verbeek FPR, Schaafsma BE, Boonstra MC, van der Vorst JR, Liefers GJ, van de Velde CJH, Frangioni JV, Vahrmeijer AL: Real-time intraoperative detection of breast cancer using near-infrared fluorescence imaging and Methylene Blue. European Journal of Surgical Oncology (EJSO) 2014, 40(7):850–858.

33. Matsui A, Tanaka E, Choi HS, Kianzad V, Gioux S, Lomnes SJ, Frangioni JV: Real-time, near-infrared, fluorescence-guided identification of the ureters using methylene blue. Surgery 2010, 148(1):78–86.

34. DiSanto AR, Wagner JG: Pharmacokinetics of highly ionized drugs. I. Methylene blue--whole blood, urine, and tissue assays. J Pharm Sci 1972, 61(4):598–602.

35. DiSanto AR, Wagner JG: Pharmacokinetics of highly ionized drugs. II. Methylene blue--absorption, metabolism, and excretion in man and dog after oral administration. J Pharm Sci 1972, 61(7):1086–1090.

36. Datta S, Wheatstone S, Challacombe B: The acute management of iatrogenic urological injuries; strategies and mind-set for the urologist attending an unfamiliar operating theatre. BJU Int 2013, 112(5):540–542.

37. Brandes S, Coburn M, Armenakas N, McAninch J: Diagnosis and management of ureteric injury: an evidence-based analysis. BJU Int 2004, 94(3):277–289.

38. Delacroix SE, Jr., Winters JC: Urinary tract injuries: recognition and management. Clin Colon Rectal Surg 2010, 23(3):221.

39. Yu M, Liu J, Ning X, Zheng J: High-contrast Noninvasive Imaging of Kidney Clearance Kinetics Enabled by Renal Clearable Nanofluorophores. Angew Chem Int Ed Engl 2015, 54(51):15434–15438.

40. Yu M, Zhou J, Du B, Ning X, Authement C, Gandee L, Kapur P, Hsieh JT, Zheng J: Noninvasive Staging of Kidney Dysfunction Enabled by Renal-Clearable Luminescent Gold Nanoparticles. Angew Chem Int Ed Engl 2016, 55(8):2787–2791.

41. Huang J, Lyu Y, Li J, Cheng P, Jiang Y, Pu K: A Renal-Clearable Duplex Optical Reporter for Real-Time Imaging of Contrast-Induced Acute Kidney Injury. Angew Chem Int Ed Engl 2019, 58(49):17796–17804.

42. Huang J, Xie C, Zhang X, Jiang Y, Li J, Fan Q, Pu K: Renal-clearable Molecular Semiconductor for Second Near-Infrared Fluorescence Imaging of Kidney Dysfunction. Angew Chem Int Ed Engl 2019, 58(42):15120–15127.

43. Huang J, Weinfurter S, Daniele C, Perciaccante R, Federica R, Della Ciana L, Pill J, Gretz N: Zwitterionic near infrared fluorescent agents for noninvasive real-time transcutaneous assessment of kidney function. Chem Sci 2017, 8(4):2652–2660.

44. Antaris AL, Chen H, Cheng K, Sun Y, Hong G, Qu C, Diao S, Deng Z, Hu X, Zhang B, Zhang X, Yaghi OK, Alamparambil ZR, Hong X, Cheng Z, Dai H: A small-molecule dye for NIR-II imaging. Nat Mater 2016, 15(2):235–242.

45. Semonin OE, Johnson JC, Luther JM, Midgett AG, Nozik AJ, Beard MC: Absolute Photoluminescence Quantum Yields of IR-26 Dye, PbS, and PbSe Quantum Dots. Jphyschemlett 2010, 1(16):2445–2450.

