## Supporting information for "Structural and Functional NIR-II Fluorescence Bioimaging in Urinary System via Clinically Approved Dye Methylene Blue"

D. Xue, Z. Lu, A. Zebiluba, Prof. J. Qian and Prof. G. Li

Department of Urology, Sir Run Run Shaw Hospital, School of Medicine, Zhejiang University

Hangzhou 310016, China

Z. Feng and Prof. J. Qian

State Key Laboratory of Modern Optical Instrumentations, Centre for Optical and Electromagnetic Research, College of Optical Science and Engineering, Zhejiang University

Hangzhou, 310058, China.

D. Wu

Department of General Surgery, Sir Run Run Shaw Hospital, School of Medicine, Zhejiang University

Hangzhou, 310016, China.

**Abbreviations**

NIR: near-infrared; MB: methylene blue; FDA: Food and Drug Administration; InGaAs: indium gallium arsenide; QY: quantum yield; FWHM: full-width-half-maximum; SBR: signal to background ratio; PBS: phosphate buffer saline; DI: deionized; UUO: unilateral ureteral obstruction; URF: unilateral renal failure. TFICs: time-fluorescence intensity curves; DCE: Dichloroethane; LK: left kidney; RK: right kidney. BUN: urea nitrogen; Cre: serum creatinine; OD: optical density; i.v.: intravenously; b.i.: bladder irrigation

**Supplementary figures**

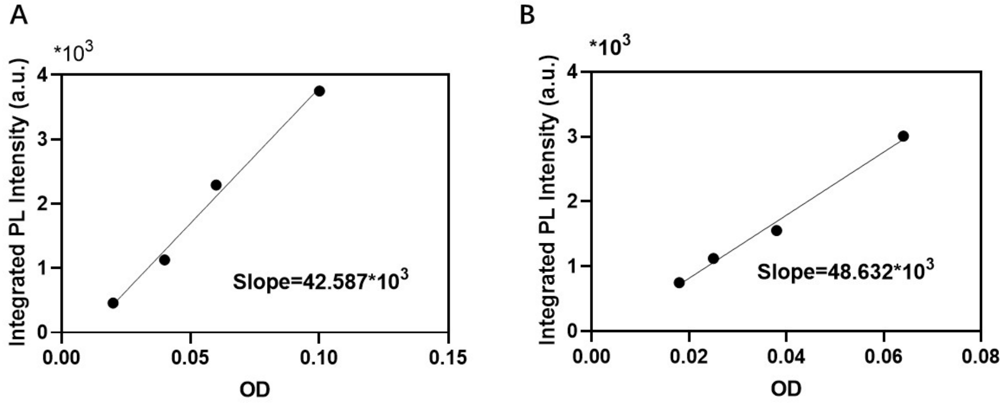


**Figure S1.** Integrated NIR-II fluorescence intensities (1000-1500 nm) plotted a function of OD at 623 nm for **(A)** MB aqueous solution and **(B)** IR-26 in DCE (reference solution). The data were fitted into linear functions with slopes of 48632 for IR-26 in DCE and 42587 for MB in water. The measured quantum yields of MB aqueous solution is 0.2%.

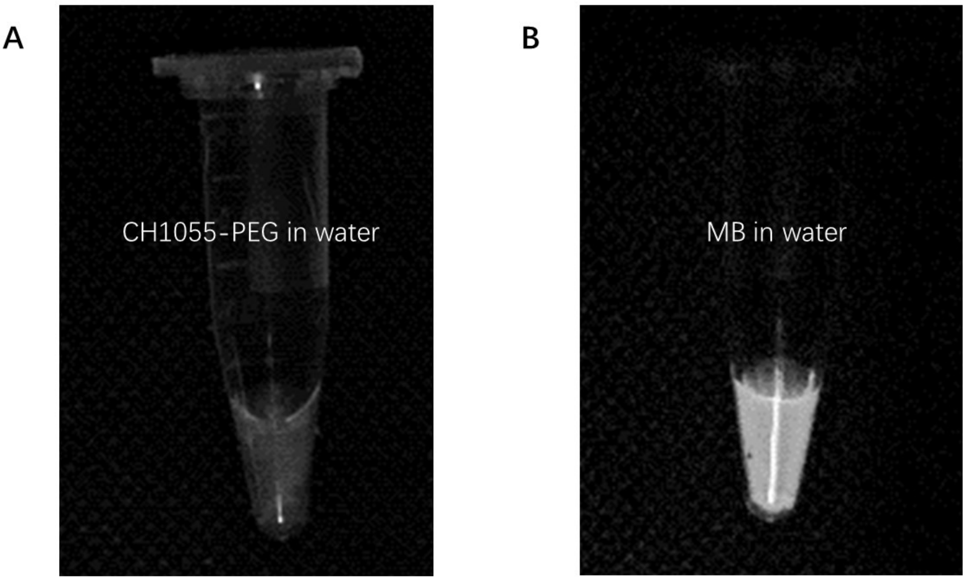


**Figure S2.** NIR-II fluorescence images of (A) CH-1055-PEG in water and (B) MB in the water at the same concentration (0.1mg/mL) under the 793 nm laser (20 mW/cm$^2$) and 623 nm LED irradiation (20 mW/cm$^2$), respectively. Exposure time: 20 ms.

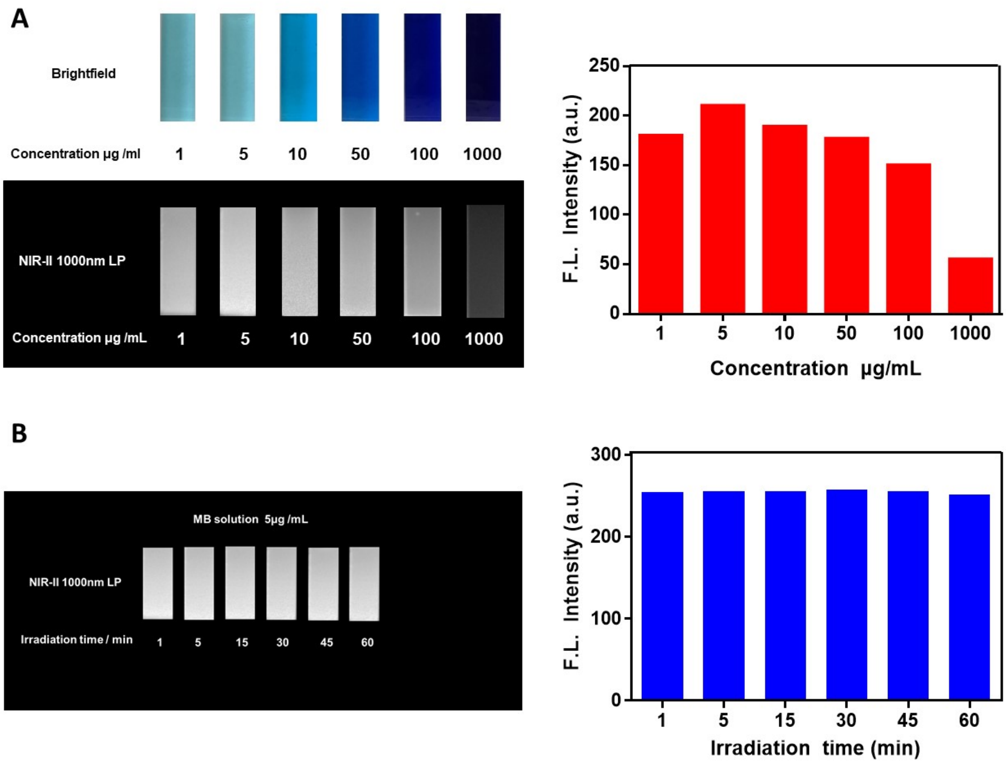


**Figure S3. (A)** NIR-II fluorescence (with a 1000-nm long-pass filter) and bright-field images of vials filled with MB aqueous solution at 6 different concentrations. (B) The photostability and NIR-II fluorescence intensity measurement of MB aqueous solution under the excitation of a 623 LED at different time points during LED illumination (density, 80 mW/cm$^2$).

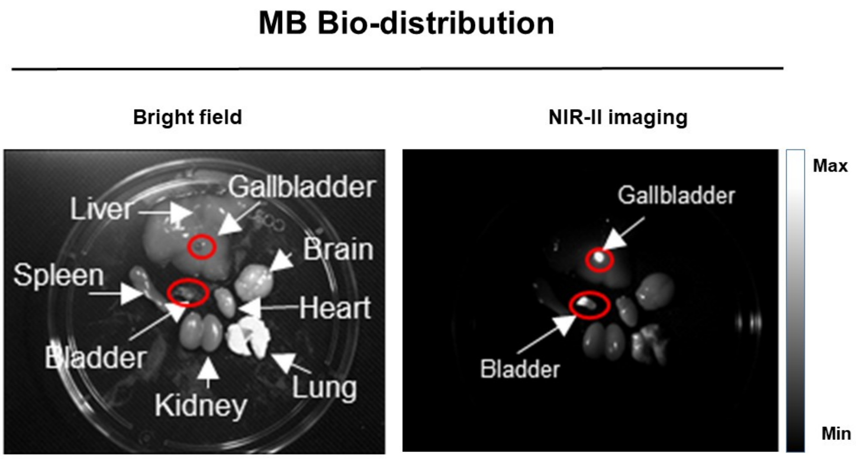


**Figure S4.** Ex vivo NIR-II fluorescence images of major organs dissection from the mouse at 1 h post the intravenous injection of MB (0.5 mg/mL, 200 μL). All images were taken under 623 nm LED excitation (50 mW/cm2). Exposure time: 30 ms.

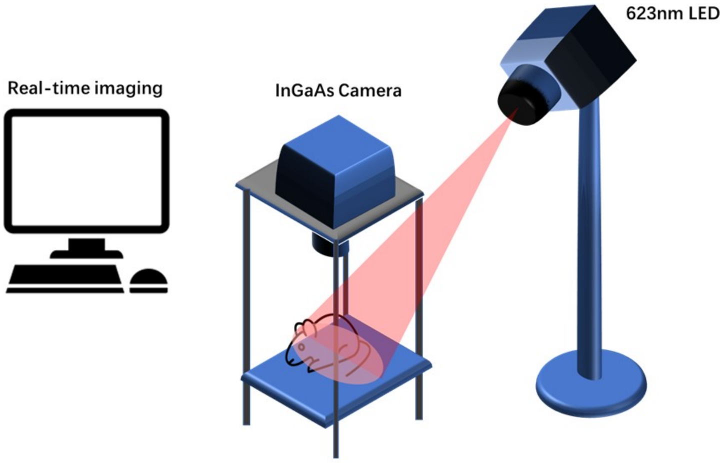


**Figure S5.** Schematic illustration for NIR-II fluorescence imaging system
